# Post-transcriptional regulation of cellulose synthase genes by small RNAs derived from CESA antisense transcripts

**DOI:** 10.1101/2020.04.30.070854

**Authors:** Daniel B. Nething, John W. Mishler-Elmore, Michael A. Held

## Abstract

Transcriptional regulatory mechanisms governing plant cell wall biosynthesis are incomplete. Expression programs that activate wall biosynthesis are well understood, but mechanisms that control the attenuation of gene expression networks remain elusive. Previous work has shown that small RNAs (sRNAs) derived from the *HvCESA*6 (*Hordeum vulgare, Hv*) antisense transcripts are naturally produced and are capable of regulating aspects of wall biosynthesis. Here, we further test the hypothesis that *CESA*-derived sRNAs generated from *CESA* antisense transcripts are involved in the regulation of cellulose and broader cell wall biosynthesis. Antisense transcripts were detected for some, but not all members of the *CESA* gene family in both barley and *Brachypodium distachyon*. Phylogenetic analysis indicates that antisense transcripts are detected for most primary cell wall CESA genes, suggesting a possible role in the transition from primary to secondary cell wall biosynthesis. Focusing on one antisense transcript, *HvCESA1* shows dynamic expression throughout development, is correlated with corresponding sRNAs over the same period and is anticorrelated with *HvCESA1* mRNA expression. To assess the broader impacts of *CESA*-derived sRNAs on the regulation of cell wall biosynthesis, transcript profiling was performed on barley tissues overexpressing *CESA*-derived sRNAs. Together the data support the hypothesis that *CESA* antisense transcripts function, through an RNA-induced silencing mechanism, to degrade *cis* transcripts, and may also trigger *trans*-acting silencing on related genes to alter the expression of cell wall gene networks.

## Introduction

As young plant cells grow and divide, they produce thin and elastic primary cell walls (PCWs). When cell growth ceases, certain cell types will undergo cell wall thickening to form rigid secondary cell walls (SCWs). The major polysaccharide for both PCW and especially SCW is cellulose. Cellulose is made by plasma membrane resident glycosyltransferases (GTs) called cellulose synthases (CESAs). CESAs synthesize individual β-(1,4) linked glucan chains, which associate to form larger paracrystalline microfibrils. Individual CESA proteins interact to form large, rosette-shaped cellulose synthase complexes (CSCs) (Brown and Montezinos 1976; Mueller and Brown 1980; Giddings et al. 1980; Herth 1985; Kimura et al. 1999). The exact number of CESA proteins in a given CSCs is unclear, but current models describe it as a hexamer of trimers that utilize at least three unique non-redundant CESA isoforms (Taylor et al. 2000; Taylor et al. 2003; Gonneau et al. 2014; Hill et al. 2014). Additionally, PCW and SCW CSCs use different sets of CESAs. In *Arabidopsis thaliana* for example, *AtCESA*s *1, 3*, and *6*/*2*/*5* are highly co-expressed and interact to form PCW CSCs (Persson et al. 2007) while *AtCESA*s *4, 7*, and *8*, are highly co-expressed and form SCW CSCs (Brown et al. 2005; Persson et al. 2005). All plants examined to date have co-expressed orthologs of each of these *Arabidopsis CESA*s indicating conservation across plant lineages (Carroll and Specht 2011). In *Hordeum vulgare* (barley) for example, *HvCESA*s *1, 2*, and *6* are co-expressed and comprise CSCs for PCWs, while *HvCESA*s *4, 7*, and *8* are for SCW CSCs (Burton et al. 2004).

PCW and SCW formation each require the concerted action of many additional GTs and cell wall modifying enzymes. Hemicellulose and pectin GTs, needed for PCW formation, tend to be co-expressed with PCW *CESA*s, while GTs and lignin biosynthetic enzymes tend to be co-expressed with SCW *CESA*s (Persson et al. 2005; Brown et al. 2005; Mutwil et al. 2009). Thus, PCWs and SCWs are each synthesized by the products of specific gene networks. Importantly, as cells begin to cease cell growth, there is a transition from PCW to SCW gene networks. The factors that drive this transition are not fully understood, but are beginning to come to light (Li et al. 2016; Watanabe et al. 2018).

As might be expected, the actions of hormones and transcription factors are major players in regulating cell wall gene networks. Auxin, abscisic acid, brassinosteroids, cytokinins, ethylene, and giberellic acid have been shown to play various roles in SCW formation (Didi et al. 2014). Transcription factor (TF) networks have been identified as activators of primary (Sakamoto et al. 2018; Saelim et al. 2019) and secondary wall biosynthetic programs both naturally and in response biotic and abiotic stresses (Kubo et al. 2005; Mitsuda et al. 2005, 2007; Zhong et al. 2006; McCarthy et al. 2009; Zhou et al. 2009; Zhong et al. 2010; Yamaguchi and Demura 2010; Wang and Dixon 2012; Ko et al. 2012, 2014; Hussey et al. 2013; Zhong and Ye 2014; Nakano et al. 2015; Zhang et al. 2018). While much is known about activation and up-regulation of cell wall synthesizing components, the corresponding mechanisms that selectively down-regulate the same gene networks are still unclear (Wang and Dixon 2012; Li et al. 2016).

Previous work has demonstrated that the transition from PCW to SCW may be regulated in part at the post-transcriptional level by *CESA*-derived small RNAs (sRNAs) (Held et al. 2008). Here, we test the hypothesis that cell wall gene networks can be regulated by antisense RNA-derived sRNAs centered around the expression of *CESA* genes. A survey of barley and *Brachypodium distachyon CESA* genes for additional antisense transcripts was performed. Antisense transcripts were detected for some, but not all *HvCESA* genes, with a concentration on PCW *CESA*s. A developmental time course of one of these antisense transcripts (*HvCESA1*) and its corresponding sRNAs over time also showed a correlated relationship. This analysis was extended to the closely related grass, *Brachypodium* to see if this phenomenon was unique to barley. Antisense RNAs were also detected for some but not all *BdCESAs*, and were generally confined to direct barley orthologs, suggesting evolutionary conservation. Lastly, cell wall gene expression profiling was performed to examine the extent to which *CESA* sRNAs can impact the expression of cell wall gene networks. The data show close and distant targeting of cell wall-related genes moderated by sRNA mechanisms demonstrating the potential for broader cell wall gene network regulation.

## Methods

### Plant growth and tissue collection

Seeds of *Hordeum vulgare* cv. black hulless were imbibed in aerated water for 24 hours to stimulate germination. Imbibed seeds were transferred to moist vermiculite and placed in the dark at 28°C until hypocotyls emerged, generally 3-5 days. Seedlings were then transferred to autoclaved soil (Promix BX) supplemented with Osmocote (Scotts) 14-14-14 slow release fertilizer (1.8 g/L). Seedlings were grown in a Percival E36HOX growth chamber under high intensity fluorescent lamps (450-700 μmol m^-2^ sec^- 1^) programmed for a 16-hour photoperiod (25 °C day, 20°C night).

*Brachypodium distachyon* seeds were imbibed in aerated water for 48 hours to stimulate germination, then transferred to damp vermiculite and incubated at 22°C in the dark for 7 days to stimulate cotyledon growth. On day 9, seedlings were transferred to autoclaved soil (Promix BX) supplemented with Osmocote (Scotts) 14-14-14 slow release fertilizer (1.8 g/L). Seedlings were grown in a Percival E36HOX growth chamber under high intensity fluorescent lamps (180-200 μmol m^-2^ sec^-1^) programmed for a 20-hour photoperiod (22 °C constant). Third-leaf tissue from ≥ 5 plants was excised, measured for length, and pooled in liquid nitrogen at 17, 19, 21, 24, and 27 days after imbibition.

### Preparation of Barley and *Brachypodium* RNA

Pooled third-leaf samples for both survey and time course experiments were pulverized using a mortar and pestle under liquid nitrogen, and then homogenized under TRIzol® reagent (Invitrogen-Thermo/Fisher). Aliquots of each RNA sample were treated for DNA contamination using the TURBO DNA-free kit (Invitrogen-Thermo/Fisher) per the manufacturer’s instructions for rigorous treatment. Each RNA sample (0.5 μg) was separated on a 0.7-1% agarose gel and visualized with ethidium bromide dye to check for RNA degradation. Gels were imaged by a Chemidoc EQ camera (BioRad) using Quantity One software (Version 4.5.2 Build 070) to verify uniform RNA loadings. Gel images were analyzed using ImageJ (Version 1.49E). All time course measurements were normalized to the RNA loading.

### Detect of antisense RNA transcripts

#### Gene-specific primer design for tagged SS-RT-PCR

Gene-specific primers (GSPs) for antisense transcript detection for *HvCESA* and *BdCESA* gene families were designed using the OligoAnalyzer 3.1 software, as described previously (Held et al. 2008). Primers were verified for specificity by BLAST analysis against either the NCBI barley or *Brachypodium* transcript library. Each primer was pairwise aligned against every member of the corresponding *CESA* gene family to ensure specificity. To improve PCR specificity and eliminate the potential for artefacts and off-target, sense-derived transcripts, the *tag1* sequence was added to the 5’ end of each barley sense GSP for cDNA synthesis (Craggs et al. 2001), while the *tag2* sequence was added to the 5’ end of each *Brachypodium* sense-GSP for cDNA synthesis.

#### Preparation of CESA Antisense cDNA for family surveys

First strand cDNAs for antisense transcripts of *Hv and Bd CESAs* were synthesized from 1.7 μg of DNase-treated total RNA extracted from barley (13 days post imbibition, dpi) and *Brachypodium* third leaves (17 dpi) using the SuperScript III First-Strand Synthesis System (Invitrogen 18080-051), using tagged sense-GSPs (**Table S1**). Control cDNAs were prepared as follows; Oligo-dT-primed (OdT) cDNA; No primer control (NPC) cDNA with the primer replaced with nuclease free water; No reverse transcriptase control (NRT) cDNA with the RT enzyme replaced with nuclease free water. cDNA reactions were then treated with RNase H to remove residual complementary RNA per the manufacturer’s protocol, and then diluted in a 1:9 ratio of cDNA with nuclease free water.

#### Amplification of antisense transcripts

*For HvCESA* antisense transcripts, first-strand cDNAs synthesized for each were amplified by PCR using the corresponding antisense GSP and the *tag1* primer. For *BdCESA* antisense transcripts, first-strand cDNAs synthesized for each were amplified by PCR using the corresponding antisense GSP and the *tag2* primer (**Table S1**). Oligo dT primed cDNA was also amplified individually with each pair of *HvCESA* sense and antisense GSPs, as controls for amplicon size and sense mRNA presence. To rule out non-specific amplification by the tag primers (Tag controls), oligo dT primed cDNAs were amplified with antisense GSPs and the *tag1* primer (for barley samples) or *tag2* primer (for *Brachypodium* samples). All PCR amplifications were assembled on ice in 25 μl reactions using 5 μl of 5X Green GoTaq buffer (Promega M3001), 0.5-1 μl of each primer (10 μM), 0.5 μl of dNTPs (10 μM each), 4 *μ*l of diluted cDNA template, and 1.25 units of GoTaq polymerase. Cycling conditions for all reactions were optimized for melting temperature and extension time (**Table S1**). Barley PCR reactions were cycled with 2 minutes of activation at 95°C, followed by 35 cycles of 95°C for 1 min, optimized annealing temperature for 1 min, and 72°C for the optimized extension time. Final elongation was 72°C for 5 minutes. *Brachypodium* PCR reactions were cycled with 2 minutes of activation at 95°C, followed by 37 cycles of 95°C for 30 s, optimized annealing temperature for 30 s, and 72°C for the optimized extension time. Final elongation was 72°C for 5 minutes. At least 3 technical replicates were performed for each antisense cDNA sample. Experiments were performed with at least three biological replications.

#### HvCESA1 antisense time course analysis

First-strand cDNAs synthesized using the *HvA1-sense-tag1* GSP, were used as templates for PCR following the same assembly as the initial detection survey. Cycling conditions for reactions using *HvA1-sense* GSP primer and *tag1* primer included 2 min of activation at 95°C, followed by 34 cycles of 95°C for 1 min, 60°C for 1 min, and 72°C for 45 sec. Final elongation was 72°C for 5 minutes. Antisense transcript cycling conditions were optimized to terminate amplifications during the mid/late-log phase so that semi-quantitative densitometry could be performed. Three replicates of equal volumes of antisense PCR products for each time point were separated by agarose gel electrophoresis. Gels were imaged by a Chemidoc EQ camera using Quantity One software (Version 4.5.2 Build 070). Gel images were analyzed using ImageJ (Version 1.49E). Background subtraction was performed with a rolling ball radius of 50.0 pixels. Densitometry was performed, and then normalized to the densitometry results from the RNA loading gel.

#### Characterization of Amplicons

Equal volumes of each PCR product for each sample and control reaction were separated by agarose gel electrophoresis with ethidium bromide staining. Antisense amplification products were excised and purified with the Zymoclean Gel DNA Recovery Kit (Zymo) and cloned into the *pGEM T-Easy* vector kit (Promega). Clones were fully sequenced and confirmed as the targeted sequence. Inclusion of *tag* sequences confirmed that cDNA samples were primed by *sense-tag1* (barley samples) or *sense-tag2* (Brachypodium samples) GSP primers and thus could only be derived from endogenous antisense transcript templates.

### Ribonuclease protection assays

#### Design of HvCESA1 RPA Probes

A 400-base pair region inside the sequence of the *HvCESA1* antisense was amplified by RT-PCR from an oligo dT primed cDNA using 5’TAAGCGCCCAGCTTTCAA and 5’ GATACCTCCAATGACCCAGAAC oligonucleotide primers and GoTaq Green polymerase (Promega). The PCR product was cloned into the *pGEM T-Easy* vector (Promega). α-^32^P-UTP (Perkin Elmer Health Sciences) radiolabeled probes were prepared from linearized plasmid templates (*SpeI* or *NcoI*) having 5’ overhangs from either T7 or SP6 RNA polymerase using the MAXIscript Kit (Ambion) to produce the *HvCESA1* antisense-targeting (466nt) and *HvCESA1* sense-targeting (506nt) riboprobes respectively. A 61-nt portion of the *HvCESA1* sense-targeting riboprobe and an 82-nt portion of the *HvCESA1* antisense-targeting riboprobe were derived from the *pGEM T-Easy* vector, so empty vector probes were similarly prepared for both as negative controls.

#### HvCESA1 Time Course RPA Assay

Ribonuclease protection assays were performed by using the Ribonuclease Protection Assay (RPA) III kit (Ambion). Labeled riboprobes were gel-purified by 5% PAGE containing 8 M urea in 1XTBE buffer per kit instructions, and hybridized with 10–20 μg total RNA from either barley, yeast, or mouse for 16–18 h at 42 °C. Reaction mixtures were digested with RNase A/T1 (1:100) for 30 min at 37 °C, then stopped with inactivation buffer (Ambion) and protected fragments were precipitated by using 10 μg yeast RNA as a carrier. The protected fragments were separated by 12.5% PAGE containing 8 M urea in 1X TBE buffer. γ-^32^ATP (Perkin Elmer Health Sciences) end-labeled Decade Marker (Ambion), prepared per manufacturer’s protocol, served as the size standard. Autoradiograms of RPA gels were uniformly scanned at 600 dpi grayscale in a lossless format. The intensity of bands in the 21-24nt range were analyzed using ImageJ (Version 1.49E).

### Custom cell wall microarray analysis

#### Viral Inoculation of Barley Plants

Plant inoculations were carried out as described previously (Holzberg et al. 2002; Held et al. 2008). Third-leaf tissues from plants visibly demonstrating photobleaching were harvested 7 to 13 days after inoculation, with maximal photobleaching at about 8 days after inoculation. Senescent tissue was trimmed from the leaf tip if present, followed by snap-freezing in liquid nitrogen. Frozen VIGS-infected tissues were pulverized using a mortar and pestle under liquid nitrogen, and then combined with TRIzol® reagent (Invitrogen, Carlsbad CA). RNA was then prepared per the TRIzol® protocol.

#### Construction of Custom Microarray

A custom, single-channel, Agilent (Santa Clara, CA) microarray based on the 8×16K architecture was designed to identify genes regulated in response to cellulose synthase silencing enriched in sequences involved in cell wall biosynthesis, stress response, and RNA regulation. Each slide contained 8 arrays, with approximately 16K probes per array (Wolber et al. 2006). A total of 3778 60-mer probes were selected from a list of candidate genes by the Agilent eArray service, with four technical replications of each probe per array. Empty vector (EV) treated samples and *HvCESA*-silencing (*HvCESA*-CR2) treated samples were prepared and pre-screened for silencing of *HvCESA6* transcript levels via qPCR prior to microarray analysis to confirm a *HvCESA* family silenced state as described earlier (Held et al. 2008).

#### Microarray Hybridization and Data Extraction

VIGS-treated barley RNA samples were verified for quality by a Bioanalyzer 2100 instrument and hybridized to the custom 8×16K microarray per the manufacturer’s protocol (Agilent). Sixteen total samples were hybridized, one per array, with 6 BSMV-EV treated samples (negative control) and 10 BSMV-*HvCESA*-CR2 treated samples. Hybridized arrays were imaged with an Agilent Technologies Scanner G2505B, and signals were extracted using the Agilent Feature Extraction Tool (Version using protocol GE1_107_Sep09).

#### Processing of Microarray Data

Extracted microarray data was processed using the limma package from Bioconductor. Backgrounds were corrected using the normexp method with a +50 offset (Ritchie et al. 2007). Arrays were normalized between each other using the quantile method. All signals within 110% of the 95^th^ percentile of the negative controls for 6 or more arrays were ignored. Signals from replicate probes for each array were then averaged and used to identify differentially expressed genes (adjusted p < 0.05).

### Collection of *BdCESA* sRNA sequences

*Brachypodium* sRNASeq dataset OBD02 (GSM1266844) (Jeong et al. 2013) hosted at mpss.danforthcenter.org was queried (Nakano et al. 2006) using selected *BdCESA* nucleotide sequences. All sRNAs matching *BdCESAs* were BLASTed against the *Brachypodium* genome to ensure specificity to only *BdCESA* genes (E-value cutoff of 1E-10), and any sequences with alternate targets were omitted.

## Results

### Antisense transcripts detected for multiple barley *CESA*s

Tagged, strand-specific RT-PCR (SS-RT-PCR) (Craggs et al. 2001; Li et al. 2013) was used to survey the barley *CESA* gene family for antisense transcripts in barley third-leaves (Burton et al. 2004; Held et al. 2008). The presence of antisense RNAs were tested for *HvCESA1* (MLOC_55153.1), *HvCESA2* (MLOC_62778; AK366571), *HvCESA4* (MLOC_66568.3), *HvCESA5/7* (MLOC_43749; AK365079), *HvCESA6* (MLOC_64555.1), and *HvCESA8* (MLOC_68431.4). *HvCESA3* (MLOC_61930.2) was omitted from this study because its expression did not cluster with either primary or secondary-wall expression (Burton et al. 2004). To ensure antisense strand specificity, a tag sequence (*tag1*) was added to the 5’ end of each barley gene-specific cDNA synthesis primer (Craggs et al. 2001) (**Fig 1A**). Antisense transcripts were detected for *HvCESA1, HvCESA4*, and *HvCESA6*, with lengths of 913, 966, and 898 nucleotides respectively (**Fig 1B**). DNA sequencing confirmed that the antisense transcripts were complementary to the corresponding exonic sequence with no introns or indels. Further, all three amplicons included the *tag1* sequence on the correct end of the transcript, confirming that the PCR product was the direct product of an antisense-transcript. Control, sense amplicons of the same sizes (minus the length of the tag) were detected for each *HvCESA*, and showed much brighter bands, despite being cycled under the same conditions, indicating that their relative quantity is very high compared to corresponding antisense transcripts. No antisense transcripts were observed for the remaining *HvCesAs* (**Fig 1B**).

**Figure 1.**
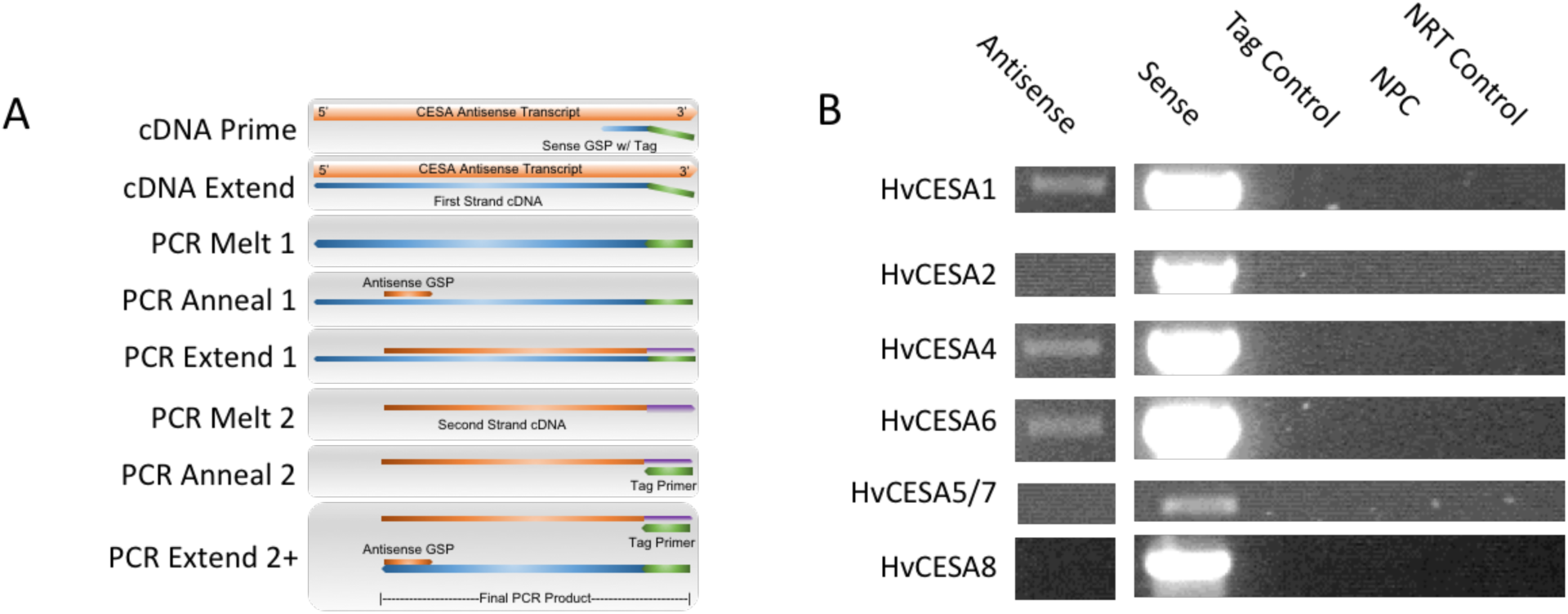
A survey of the *HvCESA* family for antisense transcripts. (A) Schematic representation of tagged, SS-RT-PCR for antisense transcript detection. First strand cDNA synthesis uses a sense gene specific primer (GSP) that is reverse-complementary only to putative antisense transcripts. To minimize PCR artifacts, a unique tag is added to the 5’ end of the sense GSP for first strand cDNA synthesis. Tagged cDNA is amplified with an antisense GSP and the tag primer. Thus, only *bona fide* antisense transcripts will be amplified. (B) Tagged, SS-RT-PCR of barley third leaf RNA for the detection of *HvCESA* antisense transcripts. PCR was performed with antisense GSPs and tag primer for Antisense, Tag control, no-primer control (NPC), and no RT (NRT) control samples. Sense transcripts were amplified using both antisense and (untagged) sense GSPs from oligo dT primed cDNA. Identity of the tagged, antisense PCR products was confirmed by DNA sequencing. See Table S1 for individual primer sequences.

### Expression of *HvCESA1* antisense and sense transcripts anticorrelate during leaf growth

*HvCesA1* antisense transcripts were monitored during barley third leaf development as previously described for *HvCESA6* (Held et al. 2008) using the tagged SS-RT-PCR method. Untagged SS-RT-PCR was used to track the *HvCesA1* sense transcript levels. The quantity of *HvCesA1* antisense transcript was lowest on day 10, then increased to a maximum on days 15 and 16 by a factor of ∼2.5-4.5 (**Fig 2B** and **Fig S1**). Over the same time period, *HvCesA1* sense signal was highest on days 10 to 13, then fell by approximately half on days 14 to 16 (**Fig 2B**). The accumulation of *HvCESA1* antisense transcripts, coupled with the decrease of *HvCESA1* sense transcripts are similar to those previously observed for *HvCesA6* (Held et al. 2008).

**Figure 2.**
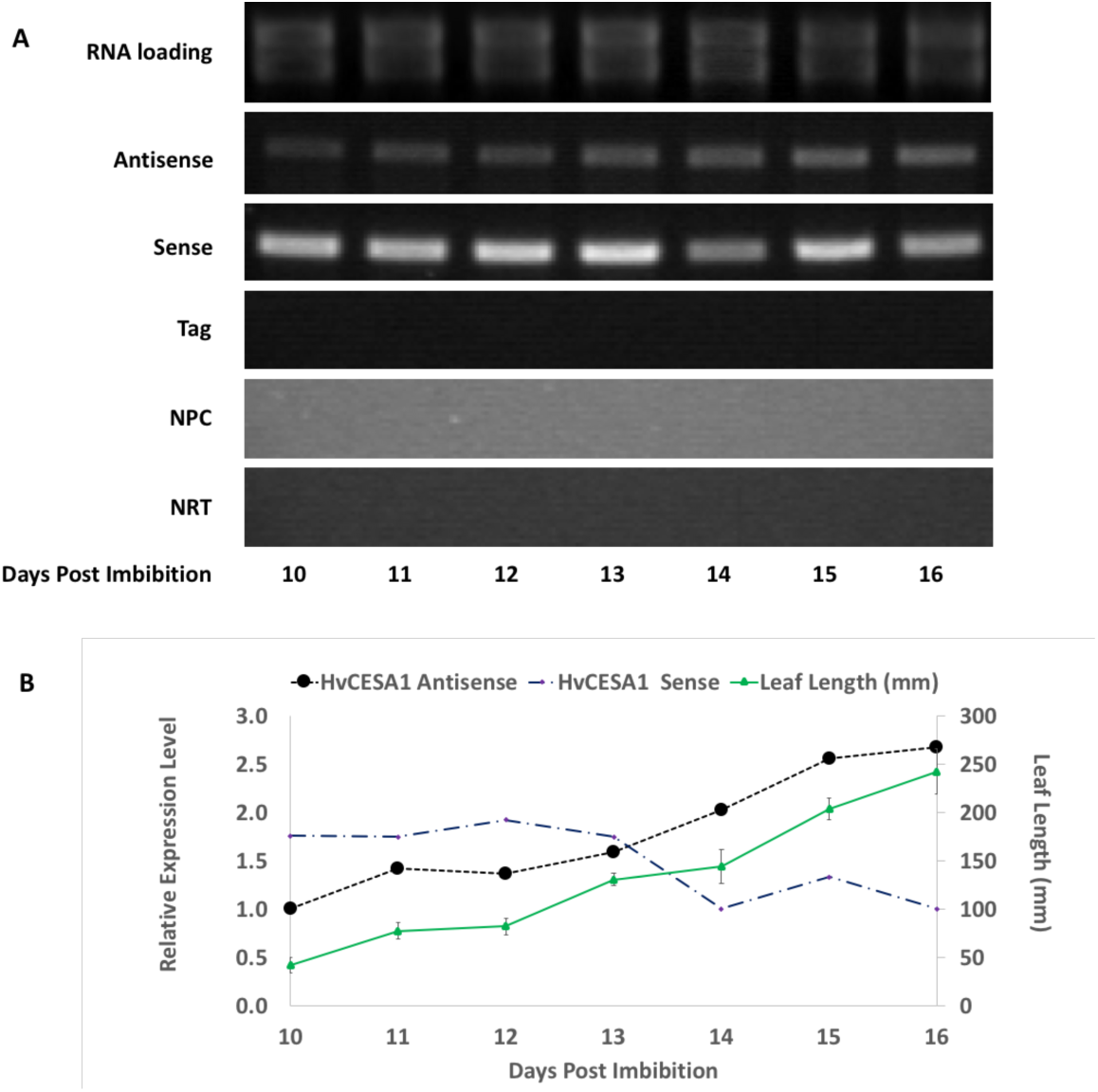
Detection of *HvCesA1* antisense transcripts by SS-RT-PCR. (A) Tagged SS-RT-PCR was performed to detect *HvCesA1* antisense transcripts over the course of third leaf development (10-16 days post imbibition). First-strand cDNAs were prepared using *HvA1-sense-Tag1* GSP (antisense; NRT control), oligo dT (sense; Tag control), or no primer at all (NPC). The *HvA1-antisense* GSP and the *Tag1* primer were used for amplification of the antisense, Tag, NPC, and NRT samples. For sense amplification, *HvA1-sense* and *HvA1-antisense* GSPs were used with *oligo dT* primed cDNAs. PCR products were confirmed by DNA sequencing. (B) Gel densitometry was performed to estimate *HvCESA1* sense and antisense transcript abundances. Data were normalized to RNA loadings and expressed relative to the first day of collection (=1). Values are representative of multiple technical replicates (n ≥ 3). Overlaid are the average leaf blade lengths (mm) ± SD (n ≥ 3).

### HvCESA1 sRNAs also accumulate over development

Ribonuclease protection assays were performed to examine the presence and abundance of *CESA*-derived sRNAs over the same time period. Antisense *HvCESA1* sRNAs (∼21-24-nucleotides) were identified via a ribonuclease protection assay (RPA) using a sense RNA riboprobe (**Fig 3; Fig S2**). The sense probe was designed to be internal to the known antisense region of *HvCESA1* (**Fig S3**), so only antisense sRNAs within the *HvCESA1* antisense transcript would be detected. The signal intensity of the *HvCESA1* sRNAs varied over time, showing an overall increase in intensity from days 11 to 16. The overall dynamic increase of the signal was by a factor of ∼2.5 for bands in the 21-24nt sRNA range (**Fig 3**), a trend similar to that of the antisense transcripts and to *HvCESA6* sRNAs previously observed (Held et al. 2008).

**Figure 3.**
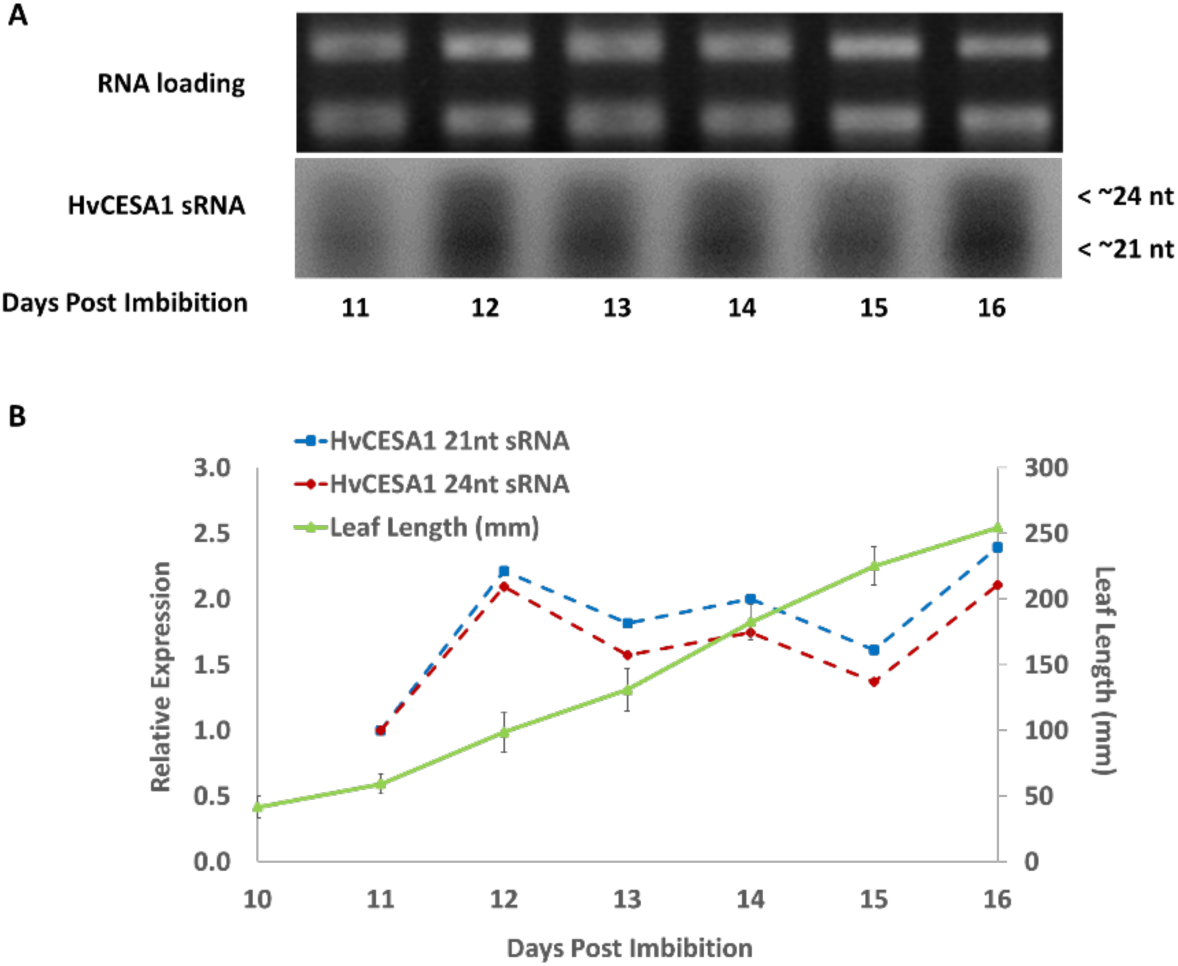
Detection of *HvCesA1* sRNAs by Ribonuclease Protection assay. (A) Ribonuclease protection assays were performed to detect *HvCESA1*-derived sRNAs across barley leaf development (11-16dpi). A sense RNA probe was used to specifically protect *HvCESA1* antisense RNAs. *HvCESA1* sRNAs (∼21-24-nts) were detected with size estimation by Decade Ladder (Ambion). (B) Densitometry was performed to evaluate the change in *HvCESA1* derived sRNA abundances. Data were normalized to RNA loadings and are expressed relative to the first day of collection (=1). Values are representative of multiple technical replicates (n ≥ 3). Overlaid are the average leaf blade lengths (mm) ± SD (n ≥ 3).

### Antisense transcripts are detected for multiple *Brachypodium CESA*s

RNA pools from rapidly growing *Brachypodium* third-leaves were assayed using tagged, SS-RT-PCR for *BdCESA1* (Bradi2g34240), *BdCESA2* (Bradi1g04597), *BdCESA4* (Bradi2g49912), *BdCESA5* (Bradi1g02510), *BdCESA6* (Bradi1g53207), *BdCESA7* (Bradi3g28350), *BdCESA8* (Bradi1g54250), and *BdCESA9* (Bradi4g30540) antisense RNA transcripts (see **Table S1** for primers). *BdCESA3* (Bradi1g29060) and *BdCESA11* (Bradi1g36740) were not examined, as they each are missing specific motifs characteristic of cellulose synthases (Handakumbura et al. 2013).

PCR amplification of the antisense cDNAs yielded antisense amplicons for *BdCESA1, BdCESA4, BdCESA6*, and *BdCESA8*, with lengths of 1059, 1078, 1107, and 1009 base-pairs respectively (**Fig 4**). Multiple sequence alignment of *Brachypodium CESA*s *1* and *8* with barley *CESA*s showed that antisense transcripts were detected for orthologous PCW *CESA*s (**Fig S4**). DNA sequencing of each antisense amplicon confirmed that all transcripts were complementary and exonic (no introns or indels), and that all four amplicons included the *tag2* primer from cDNA synthesis again indicating that SS-RT-PCR products could only have come from endogenous antisense RNA transcripts. Control sense amplicons of the same sizes were detected for each *BdCESA*s and showed much brighter bands despite being cycled under the same conditions (**Fig 4**). Similar to barley, the relative quantity of *BdCESA* antisense transcripts is low compared to the sense mRNAs. No antisense transcripts for the remaining *BdCESAs* were detected despite the presence of the control sense amplicons.

**Figure 4.**
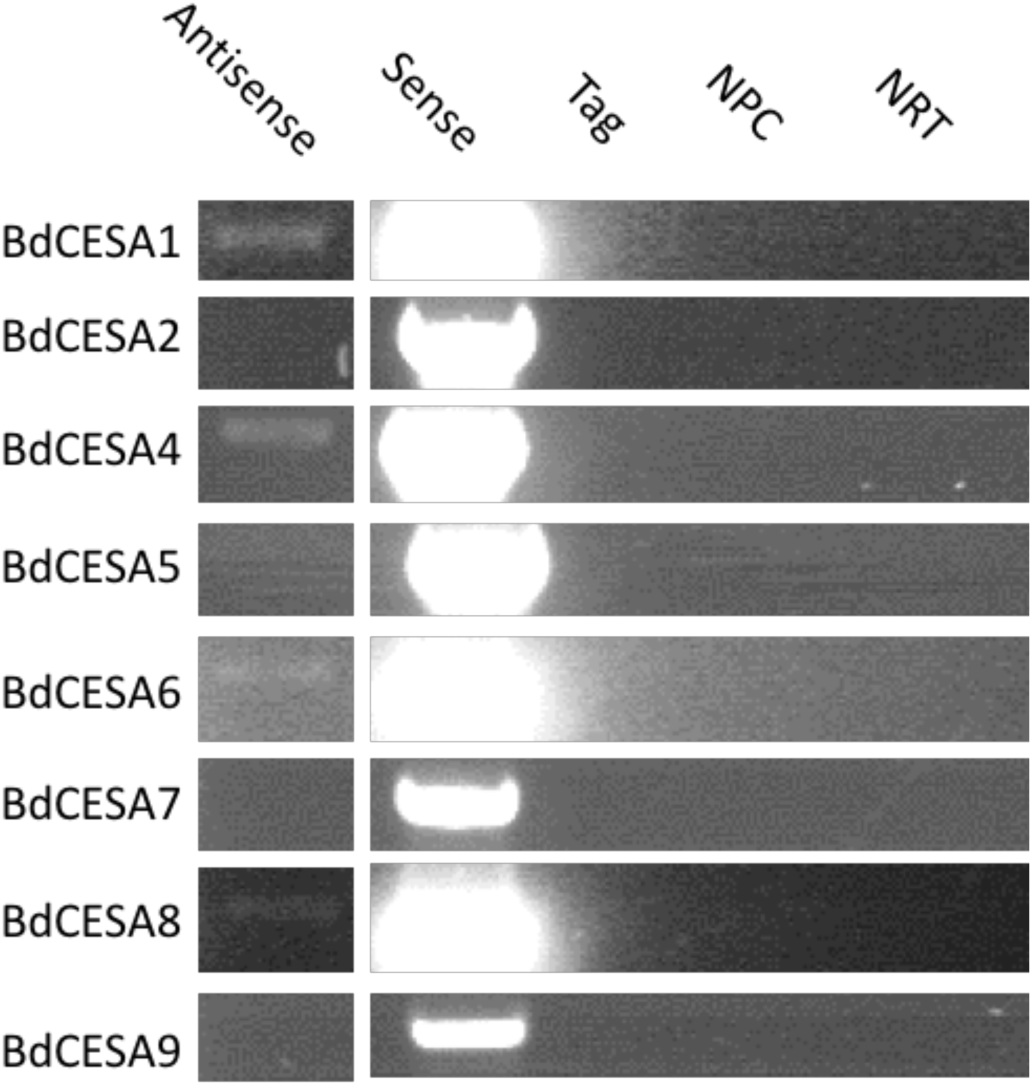
Detection of *BdCESA* antisense transcripts. Tagged SS-RT-PCR was performed to detect antisense transcripts in *Brachypodium*. First-strand cDNA was prepared using either tagged, sense GSPs for either *BdCESA*s *1, 2, 4-9* (Antisense; NRT), oligo dT primers (Sense; Tag), or no primer at all (NPC). Untagged antisense GSPs and the *tag2* primer were used for amplification of the Antisense, Tag, NPC, and NRT samples. For the sense amplification, untagged antisense GSPs were used with oligo dT primed cDNAs. PCR products were confirmed by DNA sequencing. See Table S1 for individual primer sequences.

To evaluate the presence of *BdCESA* sRNAs, sRNASeq databases were queried. Third leaf tissue data sets were not available, but similar tissue from 6-week old leaf and stem was considered comparable. sRNASeq data showed sRNA populations that matched each of the *BdCESA*s (**Table S2**). *BdCESAs* 1, 4, and 8, which produce antisense transcripts (**Fig 4**), had elevated sRNA counts compared to the other *BdCESA*s, although *BdCESA6*, which also produced antisense transcripts, had a lower count (**Table S2**). *BdCESAs* not associated with antisense transcripts, generally had lower counts, with the lone exception of *BdCESA5*. The source of *BdCESA5* derived sRNAs is unclear, but they are apparently generated independent of antisense transcripts. In general, *BdCESAs* that expressed antisense transcripts had elevated sRNA counts compared to those where antisense transcripts were not detected.

### Broad gene expression changes are observed by increasing *CESA* sRNAs

Previous work has shown silencing *HvCESA* genes by VIGS caused significant and direct reductions in *CESA* gene expression, and also caused indirect reductions in other cell wall biosynthetic genes (Held et al. 2008). That’s because VIGS of CESA genes stimulates the production of naturally abundant *CESA* sRNAs which have the potential to regulate cell wall biosynthesis in *trans*. The original study only examined a small subset of cell wall biosynthesis genes (Held et al. 2008), therefore, to more broadly examine the effects cause by over production of *HvCESA* sRNAs on cell wall gene networks, a microarray study of *CESA* VIGS-treated barley tissues was performed to compare the expression patterns of empty vector (EV) treated samples and *HvCESA*-silenced (*HvCESA*-CR2) samples. The results from the microarray indicate that 91 probes showed significant values (adj. p ≤ 0.05), with a distribution of annotated functions (**Table 1**). A total of 70 probes showed downregulated expression, while 21 probes showed upregulated expression (**Table S3**). One of the significantly down regulated genes was *HvCESA6*, a major target of the VIGS construct, confirming that silencing had indeed taken place (Held et al. 2008). Approximately 43 of the probes are specific to genes annotated for cell wall modification activity, cell wall structural proteins, glycosyltransferase activity, and glycosylhydrolase activity, suggesting the potential for broader regulatory control on cell wall gene networks via *trans* acting effects (Vasquez et al. 2004; Allen et al. 2005). If *CESA* derived sRNAs are used to help in the PCW to SCW transition, one might expect a concomitant drop in expression of genes annotated for PCW biosynthesis. While there are outliers on both sides, many down-regulated genes from this list are predicted to function in PCW biosynthesis (especially CW glycoproteins) and numerous up-regulated genes are predicted to function in SCW biosynthesis (particularly lignification) as would be expected (**Table S2**). Altogether, these data support the potential for broader cell wall gene network regulation via *CESA*-derived sRNAs.

**Table 1.** Distribution of gene annotations affected by virus-induced gene silencing (VIGS) of CESAs in barley. Protein functional groupings (protein function) are listed for genes significantly up or down regulated by CESA-VIGS as determined by microarray analysis. Number corresponds to the number of individual genes affected for each protein function category. A complete list of significantly up and down regulated genes and their functional groups is presented in Table S3.

| Protein Function | Number |
| --- | --- |
| Cell Wall Modifying Proteins | 16 |
| Transcription Factor | 16 |
| Cell Wall Structural Proteins | 12 |
| Glycosyl Transferase | 8 |
| Glycosyl Hydrolases | 7 |
| Stress Response | 6 |
| Cytoskeleton | 4 |
| Lignin Biosynthesis | 4 |
| Metabolism | 4 |
| Promoter Binding | 3 |
| Transport | 3 |
| Ribosomal | 3 |
| Epigenetic Modulator | 2 |
| Photosynthesis | 2 |
| Unknown | 1 |

## Discussion

Plant cell walls are composed of complex networks of cellulose, various hemicelluloses, pectin, lignin and glycoproteins. The amounts and proportions of these polymers vary greatly among plant cell types and across plant development. The ability of plant cells to generate wall types tailored for specific physiological roles and the ability to change wall polysaccharide biosynthesis upon various external stimuli (e.g. biotic/abiotic stresses) requires complex, multi-level regulatory control. Gene expression networks for polymer biosynthesis are co-regulated to facilitate coordinated polymer deposition, but they also need to allow flexibility to selectively respond various stresses.

Here we provide further evidence that post-transcriptional regulation is employed to selectively attenuate the expression of cellulose synthase genes and that this regulation has the potential to broadly affect the expression of other cell wall biosynthetic genes. We also show that *CESA* antisense transcripts were not restricted to barley, as they also occur in *Brachypodium*. The detection of *CESA* antisense transcripts in another plant species suggests that they might be common in all higher plants. Further, antisense transcripts were detected for several orthologous PCW *CESA*s (**Fig S4**) and therefore may represent an evolutionary conserved regulatory mechanism for limiting the expression of PCW *CESA*s.

While much is known about activation and repression of SCW gene networks, relatively little is known regarding the repression of PCW networks (Wang and Dixon 2012; Li et al. 2016). Between barley and *Brachypodium*, a total of 7 antisense transcripts were detected. Five of these antisense transcripts are produced from PCW *CESA* genes, with the lone SCW exceptions being *HvCESA4* and *BdCESA4* for barley and *Brachypodium*, respectively (**Fig S4**). While the significance of *HvCESA4* and *BdCESA4* SCW antisense transcripts are not fully understood at present, the data support our previous hypothesis that post-transcriptional sRNA regulation is important for the transition from the PCW to SCW gene network (Held et al. 2008).

Future work directed at detecting antisense transcripts in *Arabidopsis* is in progress. Moving this research into a more tractable genomic model will help shed light on the mechanisms of sRNA biogenesis. Using an inducible SCW system in *Arabidopsis* (Zuo et al. 2000; Pesquet et al. 2010) should help further clarify the roles of *CESA* sRNAs and their putative involvement in mediating the transition from PCW to SCW biogenesis.

## Acknowledgements

The authors thank the Vijayanand Nadella, William H. Broach, Rachel Yoho, and Kaiyu Shen of the Ohio University Genomics Facility for DNA sequencing, RNA bioanalyzer, and custom microarray support. Mr. A. Miner is thanked for his support during manuscript revision. This investigation was conducted in a facility constructed with support from Research Facilities Improvement Program Grant Number C06 RR-014575-01 from the National Center for Research Resources, National Institutes of Health.

## Supplemental Materials

**Table S1.**
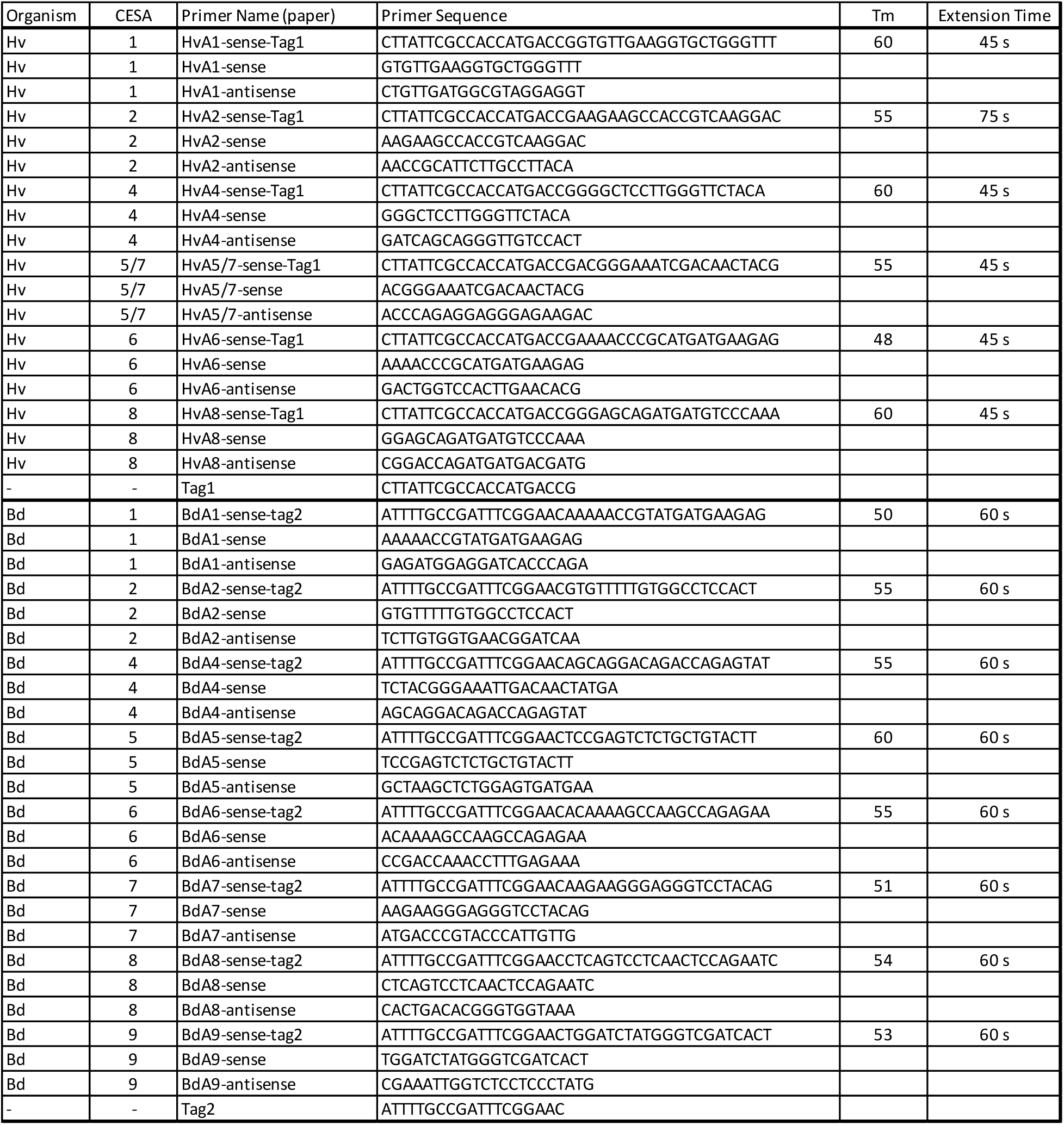
PCR primers used in this study.

**Table S2.** *BdCESA* sRNA counts mined from 6-week old stem and leaf *Brachypodium* data.

| *CESA* | Locus | Total # sRNA Reads | Unique sRNA Reads |
| --- | --- | --- | --- |
| *BdCESA1* | Bradi2g34240 | 108 | 50 |
| *BdCESA2* | Bradi1g04597 | 10 | 5 |
| *BdCESA4* | Bradi2g49912 | 90 | 44 |
| *BdCESA5* | Bradi1g02510 | 105 | 51 |
| *BdCESA6* | Bradi1g53207 | 41 | 20 |
| *BdCESA7* | Bradi3g28350 | 8 | 4 |
| *BdCESA8* | Bradi1g54250 | 79 | 38 |
| *BdCESA9* | Bradi4g30540 | 38 | 19 |

**Table S3.**
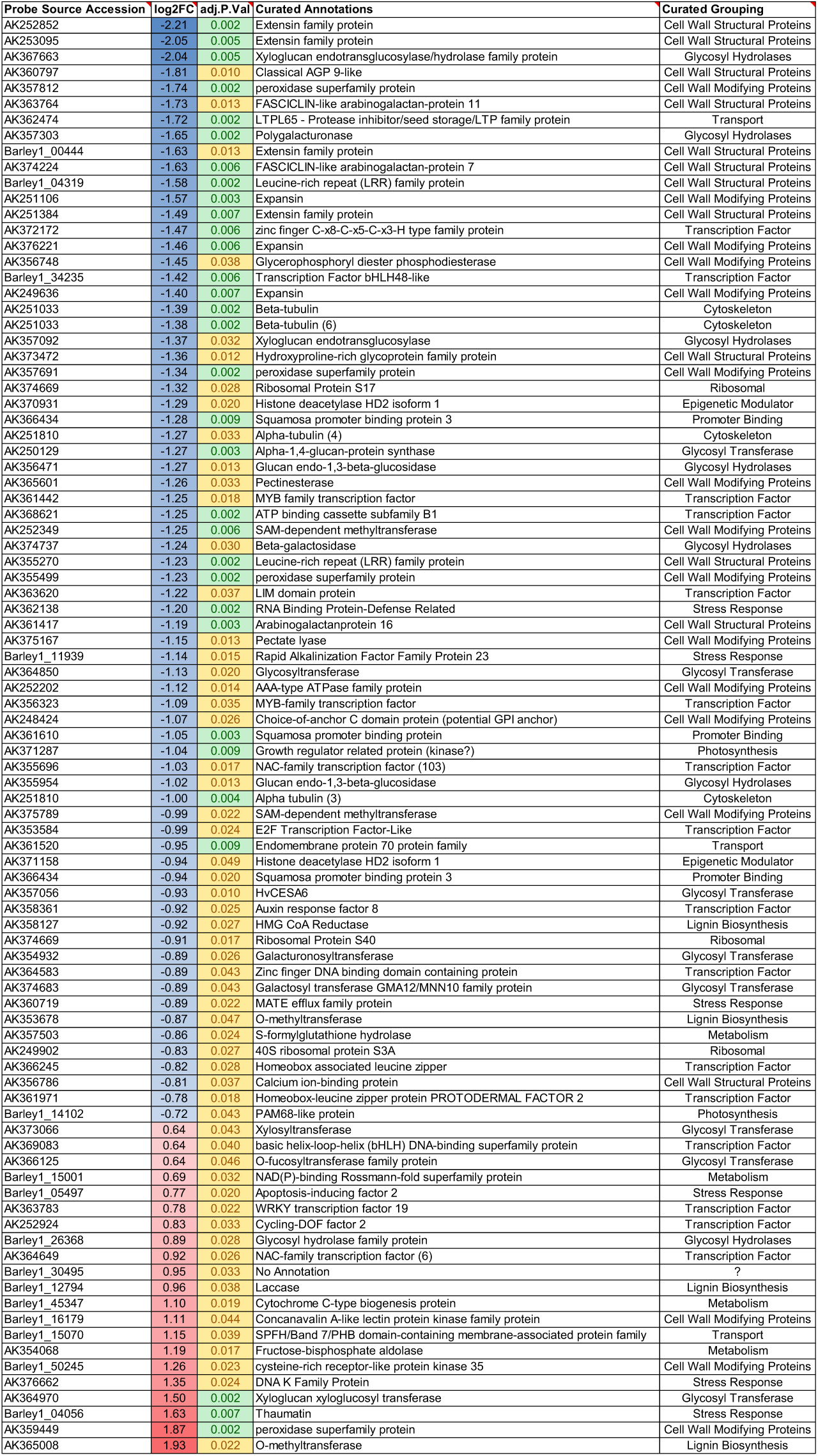
Changes in barley cell wall gene expression when the cellulose synthase gene family is specifically targeted by VIGS. Differentially expressed genes are sorted by log2 fold change.

**Figure S1.**
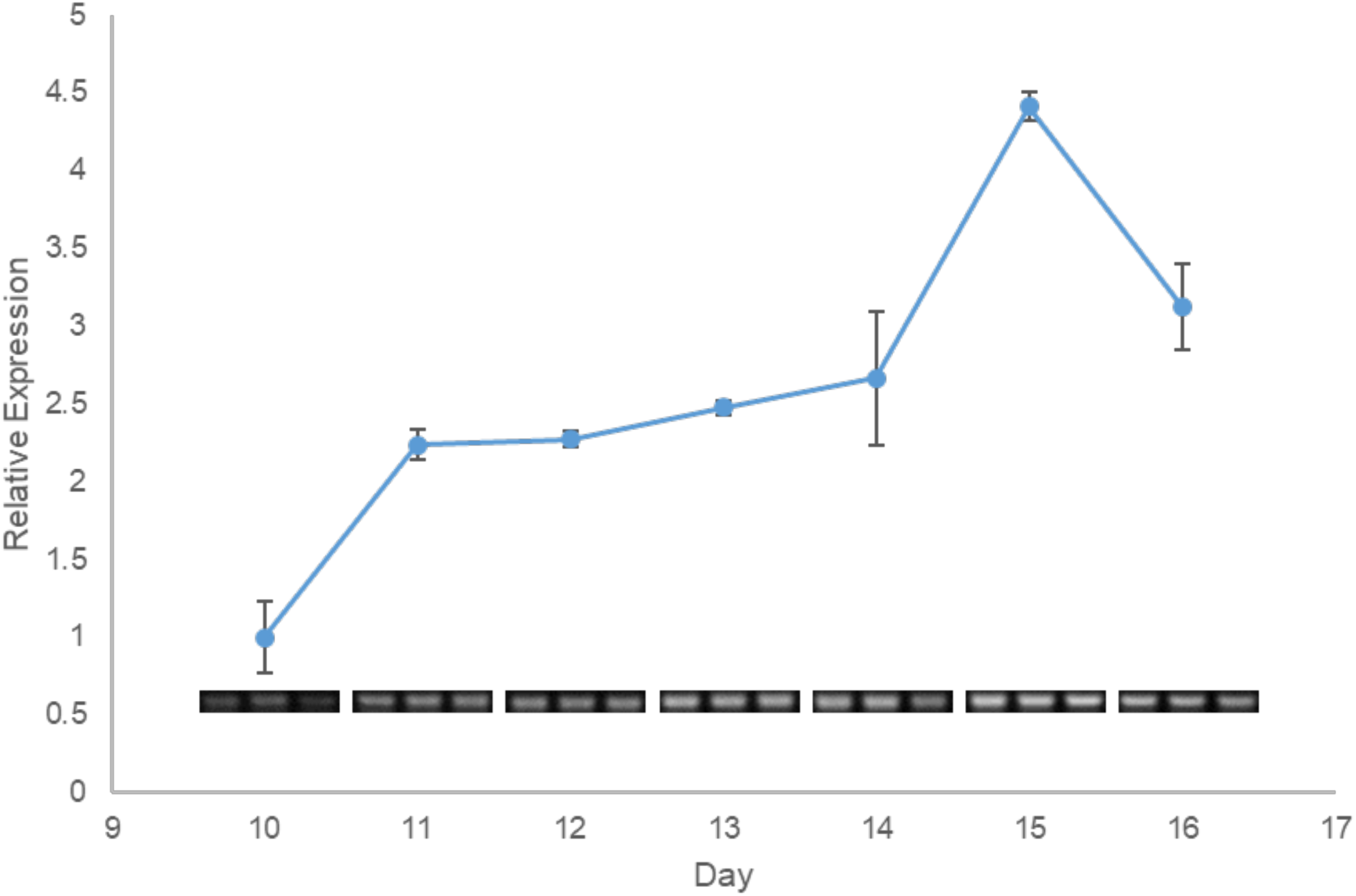
Additional time course study for *HvCESA1* antisense. SS-RT-PCR was performed to determine the expression of *HvCESA1* antisense transcripts over the course of barley third leaf development. Band intensities after agarose gel electrophoresis were determined by densitometry and expressed relative to day 10 of the time course. Values are averages of three technical replicates (gel images shown below each time point). Error bars represent SD.

**Figure S2.**
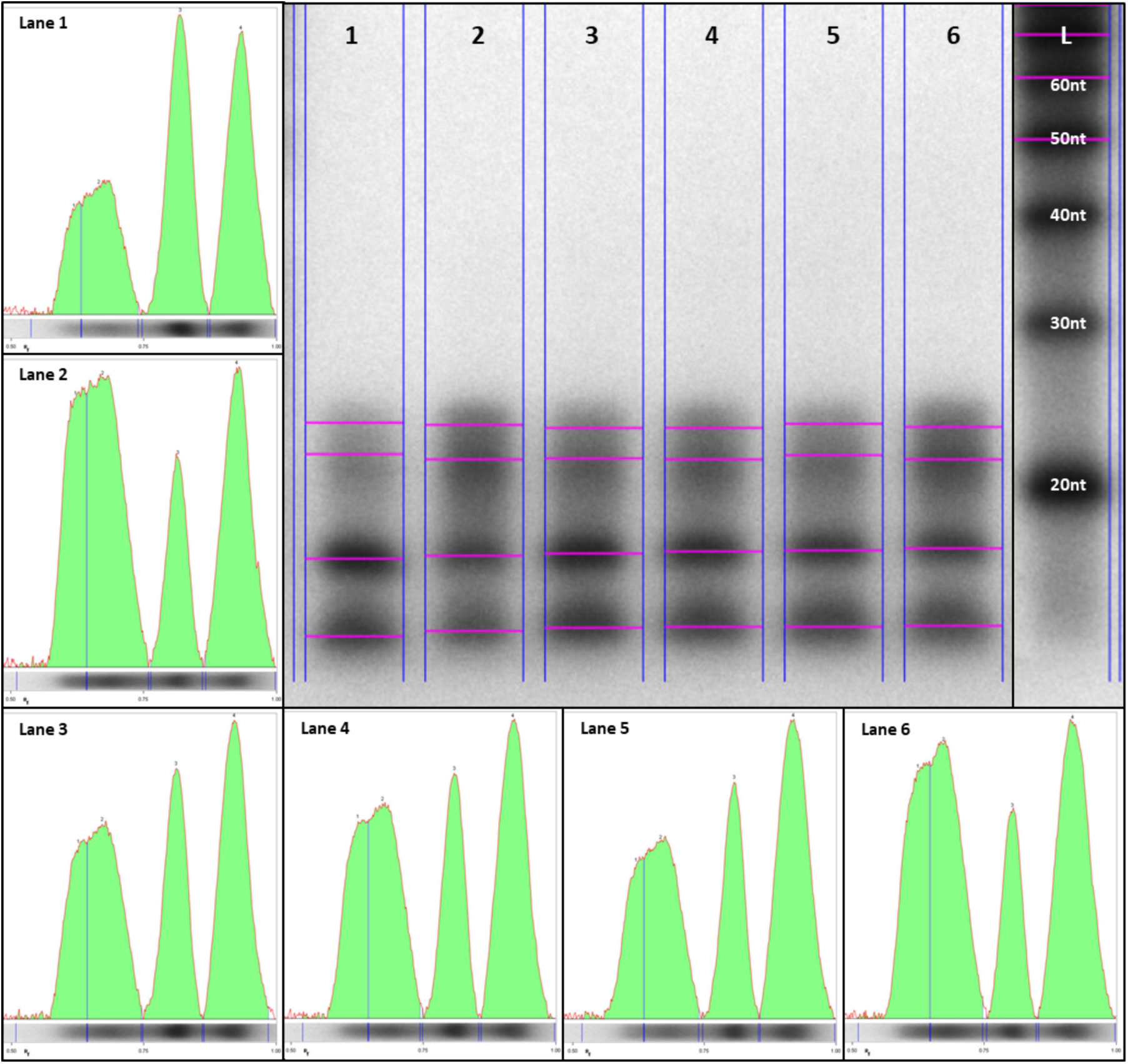
HvCESA1-associated sRNAs. Ribonuclease protection assays were performed to detect *HvCESA1*-derived sRNAs across barley leaf development 11-16dpi (Lanes 1-6). A sense RNA probe was used to protect *HvCESA1* antisense RNAs. Band intensities for lanes 1-6 are shown in individual panels to evaluate the resolution and abundance of 21-24nt bands. Smaller bands (<20 nt) are non-specific digest products common to all samples. *HvCESA1* sRNAs sizes (∼21-24nt) were estimated using γ-^32^ATP end-labeled Decade Markers (Lane L).

**Figure S3.**
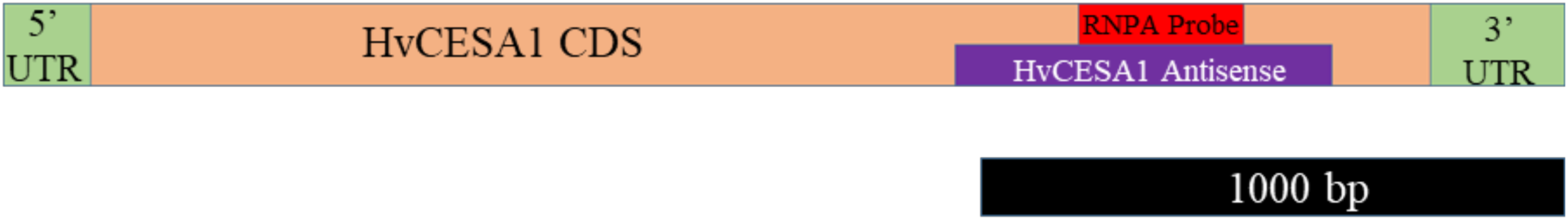
Map to scale of *HvCESA1* RPA probe and *HvCESA1* antisense transcripts. PCR amplicons and RPA probes were designed internal to the coding region of *HvCESA1*. The untranslated regions (UTR) at the 5’ and 3’ ends are indicated in green, with the coding sequence (CDS) indicated in tan. The region amplified to detect *HvCESA1* antisense transcript is in purple, and the sequence region used to probe for antisense *HvCESA1* sRNAs is indicated in red.

**Figure S4.**
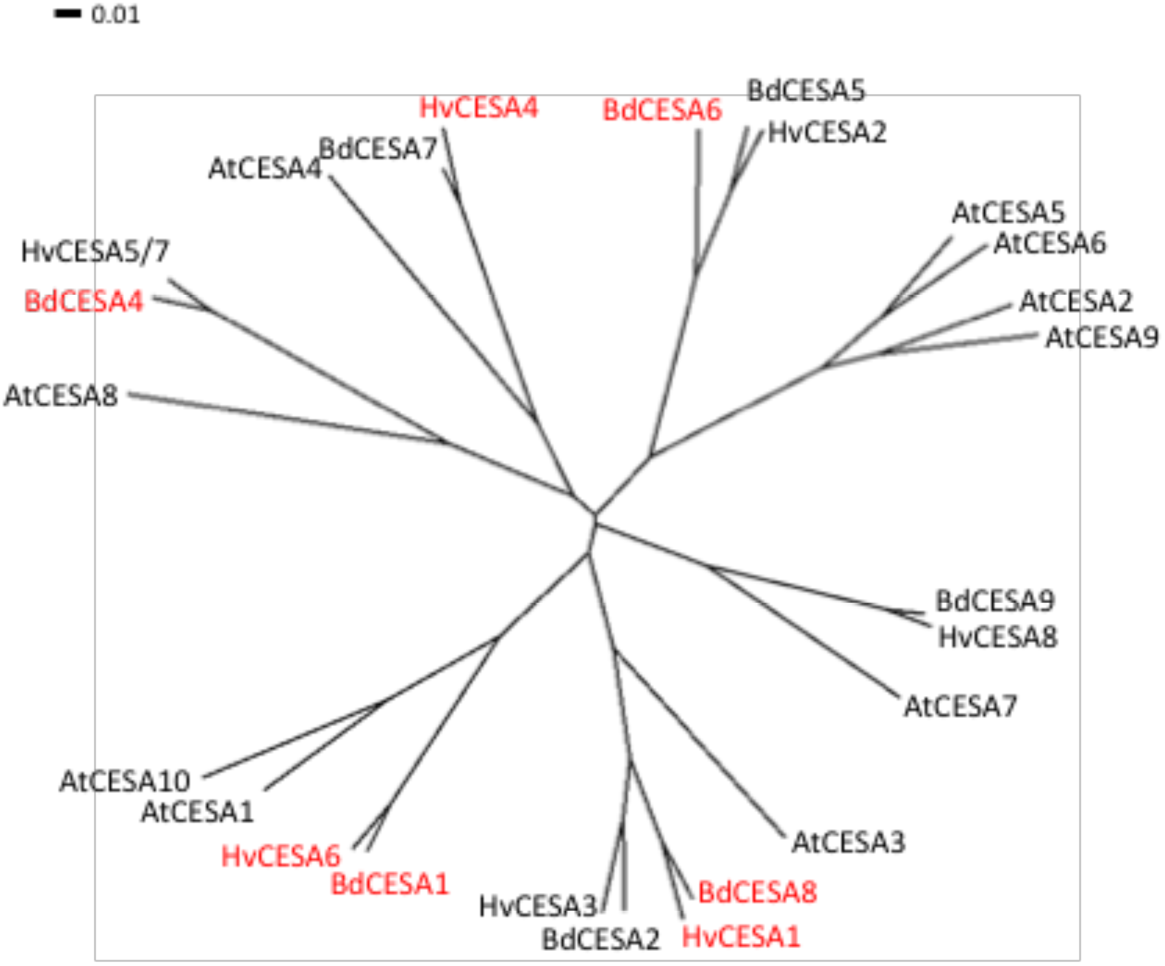
CESA Phylogenetic Tree. Multiple sequence alignment of Arabidopsis (At), barley (Hv), and *Brachypodium* (Bd) CESA proteins was performed using the Clustal Omega tool with default settings. Alignment file was loaded into Dendroscope 3 to generate an unrooted radial dendrogram. CESAs expressing antisense transcripts are highlighted in red. Protein sequences for AtCESA1 (At4g32410), AtCESA2 (At4g39350), AtCESA3 (At5g05170), AtCESA4 (At5g44030), AtCESA5 (At5g09870), AtCESA6 (At5g64740), AtCESA7 (At5g17420), AtCESA8 (At4g18780), AtCESA9 (At2g21770), and AtCESA10 (At2g25540) were collected from TAIR (https://www.arabidopsis.org/). Protein sequences for HvCESA1 (MLOC_55153.1), HvCESA2 (AK366571), HvCESA3 (MLOC_61930.2), HvCESA4 (MLOC_66568.3), HvCESA5/7 (AK365079), HvCESA6 (MLOC_64555.1), HvCESA8 (MLOC_68431.4), BdCESA1 (Bradi2g34240), BdCESA2 (Bradi1g04597), BdCESA4 (Bradi2g49912), BdCESA5 (Bradi1g02510), BdCESA6 (Bradi1g53207), BdCESA7 (Bradi3g28350), BdCESA8 (Bradi1g54250), and BdCESA9 (Bradi4g30540) were collected from PGSB (http://pgsb.helmholtz-muenchen.de/plant/barley/index.jsp).

